# Short-term transcutaneous vagus nerve stimulation increases pupil size but does not affect EEG alpha power: a replication

**DOI:** 10.1101/2023.03.08.531479

**Authors:** Beth Lloyd, Franz Wurm, Roy de Kleijn, Sander Nieuwenhuis

**Affiliations:** Institute of Psychology, Leiden University, Leiden, the Netherlands; Leiden Institute for Brain and Cognition, Leiden University, Leiden, the Netherlands

## Abstract

**Background:** Transcutaneous auricular vagus nerve stimulation (taVNS) is a promising brain stimulation method for the treatment of pharmaco-resistant epilepsy and depression. Its clinical efficacy is thought to depend on taVNS-induced activation of the locus coeruleus. However, unlike for invasive VNS, there is little evidence for an effect of taVNS on noradrenergic activity.

**Objective:** We attempted to replicate recently published findings by Sharon et al. (2021), showing that short bursts of taVNS transiently increased pupil size and decreased EEG alpha power, two correlates of central noradrenergic activity.

**Methods:** Following the original study, we used a single-blind, sham-controlled, randomized cross-over design. We applied short-term (3.4 s) taVNS in healthy human volunteers (n=29), while collecting resting-state pupil-size and EEG data. To analyze the data, we used scripts provided by Sharon and colleagues.

**Results:** Consistent with Sharon et al. (2021), pupil dilation was significantly larger during taVNS than during sham stimulation (*p* = .009; Bayes factor supporting the difference = 7.45). However, we failed to replicate the effect of taVNS on EEG alpha power (*p* = .37); the data were four times more likely under the null hypothesis (BF_10_ = 0.28).

**Conclusion:** Our findings support the effectiveness of short-term taVNS in inducing transient pupil dilation, a correlate of phasic noradrenergic activity. However, we failed to replicate the recent finding by Sharon et al. (2021) that taVNS attenuates EEG alpha activity. Overall, this study highlights the need for continued research on the neural mechanisms underlying taVNS efficacy and its potential as a treatment option for pharmaco-resistant conditions. It also highlights the need for direct replications of influential taVNS studies.

## Introduction

Vagus nerve stimulation (VNS) is an invasive technique used to electrically stimulate the vagus nerve, which has shown promise in treating pharmaco-resistant epilepsy and depression [1,2]. The vagus nerve has major afferent connections to the nucleus of the solitary tract [3], which in turn directly and indirectly modulates the locus coeruleus, the main source of norepinephrine in the central nervous system [4]. Consequently, animal studies have found that VNS increases the activity of the noradrenergic system [5–9]. Recently, transcutaneous auricular vagus nerve stimulation (taVNS) has gained traction as a non-invasive alternative to VNS [10]. taVNS is thought to mimic the effects of VNS by delivering alternating currents to the auricular branch of the vagus nerve via surface skin electrodes on the outer (left) ear. However, it is still uncertain if taVNS enhances noradrenergic activity, and hence the working mechanisms underlying taVNS are poorly understood [11].

Several studies have examined the effects of taVNS on noninvasive markers of noradrenergic activity. Invasive VNS in human patients has been found to modulate the amplitude of the P300 component of the event-related potential [12,13], a purported correlate of phasic noradrenergic activity [14,15]. However, taVNS studies have produced mixed evidence for an effect on P300 amplitude [16–19]. Studies on the effect of taVNS on salivary alpha-amylase, an indirect hormonal marker of noradrenergic activity [20,21], have also yielded mixed evidence, although a recent pooled mega-analysis found a small but significant taVNS-induced increase in salivary alpha-amylase [22]. Pupil size is considered the most reliable noninvasive marker of noradrenergic activity [23], and is consistently increased by VNS in humans [24] and animals [25,26]. So when studies began reporting null effects of taVNS on event-related pupil dilation and baseline pupil size [11,19,27–30], this led to serious doubt about the efficacy and working mechanisms of taVNS [11]. However, then an important study appeared [31], which found that short-term bursts of taVNS (3.4 s) evoked immediate but transient modulations of pupil size and EEG alpha power (8 - 12 Hz), another potential correlate of noradrenergic activity [32].

Several authors have pointed out that Sharon et al. [31] used higher stimulation intensities (individually calibrated, 2.20 ± 0.24 mA) and shorter stimulation epochs (3.4 s) than the studies cited above, most of which used a fixed intensity of 0.5 mA and stimulation epochs on the order of minutes (sometimes with a 30-s on/30-s off schedule, the regime for therapeutic use of the NEMOS® device). Therefore, the findings of Sharon and colleagues may offer important clues regarding the taVNS parameter values that are most effective in experimental research and clinical settings. However, given the many studies with null effects of taVNS, it remains to be determined if the pupil and EEG results of Sharon and colleagues are replicable. To that end, we closely replicated their experiment and systematically examined the simultaneously acquired EEG and pupil-size measurements from a sample of healthy males and females (n=29).

## Methods

### Participants

Thirty-three young adults participated in return for 17 euros or course credits. Data from four participants were excluded because of technical issues, resulting in a final sample of 29 participants (21 female; mean age 22.5 ± 3.8 years). For the EEG analysis, a further five participants were excluded due to a lack of alpha spectrum and topography in the break data. Therefore, 24 participants remained for the EEG analysis (18 female; mean age 22.2 ± 3.7 years). Individuals with neurological or cardiac disorders and individuals who received taVNS stimulation in the four weeks prior to the study were excluded from participation. Participants were instructed not to consume alcohol and caffeine in the 24 hours and three hours, respectively, before the start of the two-hour session. The study was approved by the Psychology Research Ethics Committee at Leiden University (S.T.-V1-3624).

### Procedure

The current study was carried out as closely as possible to the study of Sharon et al. [31], using task and analysis scripts kindly provided by Sharon and collagues^1^. The study design was single-blind and sham-controlled and the order of conditions (taVNS vs. sham; Figure 1A) was counter-balanced across participants. After setting up the EEG, we carried out a methods-of-limits procedure for each of the two conditions to identify the maximum comfortable stimulation level for each individual. We repeatedly applied 5 seconds of stimulation, after which participants rated the subjective intensity on a scale of 0-10 (0 = no sensation, 3 = light tingling, 6 = strong tingling, 10 = painful). The level of stimulation started at 0.1 mA and increased by 0.2 mA until a rating of 9 or a maximum stimulation level of 5 mA was reached. The intensity corresponding to the subjective rating of 9 (just below painful) was selected.

**Figure 1.**
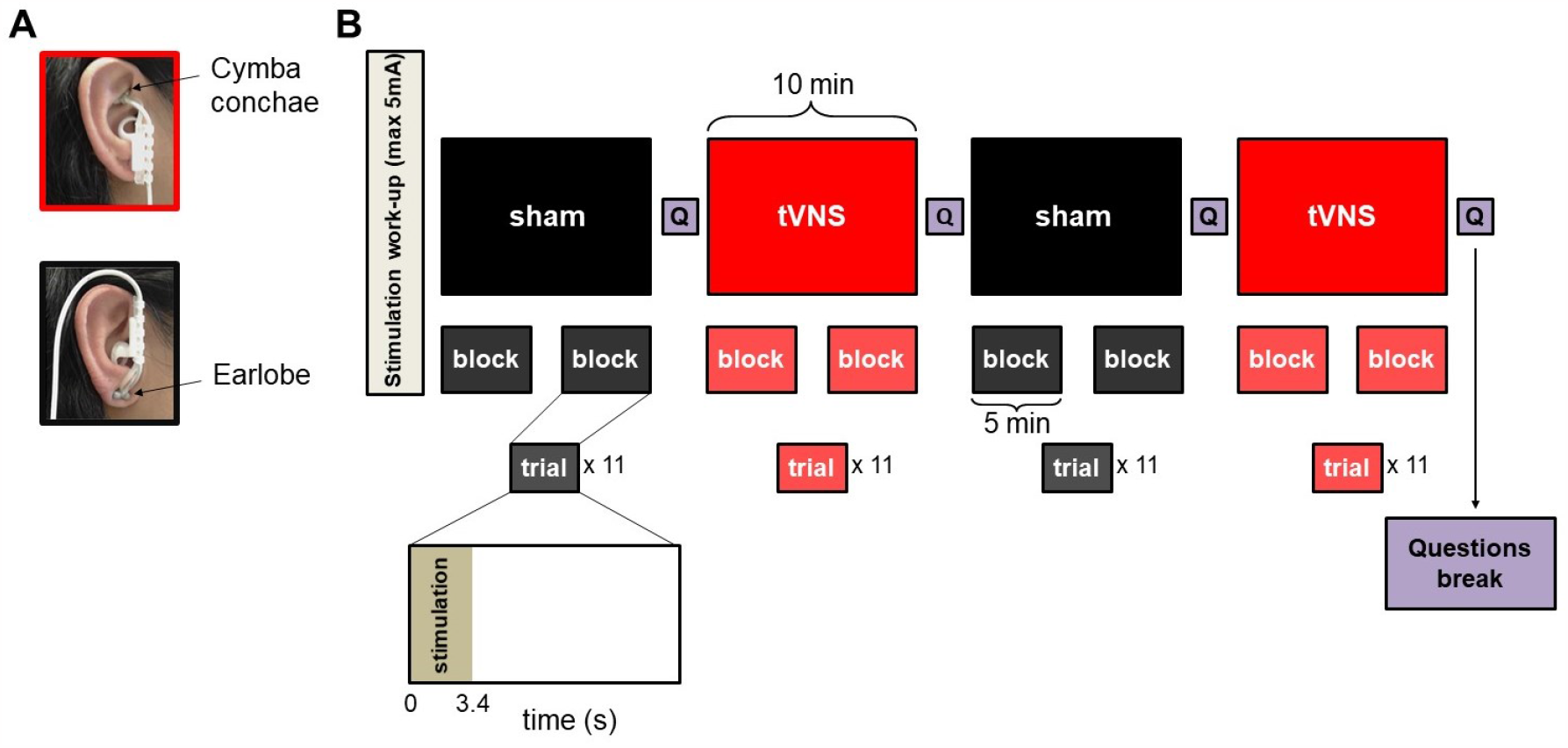
Study design for testing the effects of taVNS on pupil diameter and alpha activity. A) The placement of the electrodes at the (left) cymba conchae (taVNS condition) and the (left) earlobe (sham condition; picture adapted from [34]). B) The experimental sequence, as in [31].

Next, the participants were instructed to fixate on a white fixation cross on a grey background (RGB: 125, 125, 125). There were eight blocks of 11 trials, each lasting ∼5 minutes (Figure 1B). Each trial started with 3.4 s of stimulation, followed by an inter-stimulus interval jittered between 25 s and 27 s. After every two blocks participants responded to 11 questions regarding their subjective experience of stimulation and were free to take a break. Then the condition switched between taVNS and sham. The EEG data collected during the breaks were used to characterize individual alpha activity. At the end of the session we asked participants whether they thought the final block involved taVNS or sham stimulation. Two participants did not respond to this question. Of the other participants, 48% responded correctly, suggesting that participants could not distinguish between taVNS and sham.

### Transcutaneous auricular vagus nerve stimulation

We used the NEMOS® device (tVNS Technologies GmbH) to deliver electrical stimulation to the left cymba conchae or the left ear lobe (Figure 1A). The electrodes were covered with cotton rings soaked in electrolyte conductive liquid to establish good connection for stimulation and to prevent direct contact with the skin. Pulses (width: 200–300 µs) were delivered at a rate of 25 Hz and an intensity as described above. To achieve precisely 3.4 s of stimulation, the NEMOS® device was controlled using two linear actuators which pressed the ON/OFF buttons. These ‘push’ commands were sent at pre-programmed times via an Arduino mini, and were embedded within the task script, which was programmed using the Expyriment Python package [33]. Two checks were implemented to ensure that only trials with good connection between the electrodes and the skin were included. First, the taVNS device automatically stops stimulating once connection has been disrupted. Second, we placed an additional external EEG electrode at the back of the left earlobe, and one on the auricle. Signal from these two electrodes showed a clear artefact during stimulation. Trials in which this artefact was not visible or did not show the pattern of a steady ramp-up of stimulation with 25-Hz pulses were removed from analysis (see EEG section for details).

### Pupillometry

#### Data acquisition

The eye-tracker was positioned 75 cm from the participant’s eyes. Pupil size was recorded from the dominant eye at a sampling rate of 40 Hz using a Tobii Pro eye-tracker. Eye gaze was measured at a sampling rate of 120 Hz. Eye positions were transformed to degrees of visual angle based on a five-point calibration procedure. The experiment was carried out under constant ambient light.

#### Data analysis

Pupil data were preprocessed using PupCor (https://github.com/lindvoo/PupCor). Blinks were removed by linearly interpolating over the periods with invalid samples (automatically marked by the device manufacturer) from 100 ms before blink onset to 400 ms after blink offset. Next, the data were manually checked and corrected if any artifacts had not been successfully removed. Pupil-size data were then low-pass filtered using a 10-Hz fourth-order Butterworth filter with zero phase shift, and segmented into trials by extracting the samples from -10 s to +13.4 s around stimulation onset. Trials in which > 50% of the samples were marked as invalid were excluded, resulting in an average of 38.9 ± 10.7 trials remaining for the taVNS condition and 39.1 ± 10.5 trials for the sham condition. As in Sharon et al. [31], pupil size was converted to ‘percentage change’ values relative to a 10-s baseline before stimulation onset using the formula: 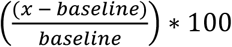. Baseline pupil size did not differ between conditions (*M*_*taVNS*_ = 3.69 ± 0.6 mm, *M*_*sham*_ = 3.73 ± 0.72 mm, *p* = .59).

After this preprocessing procedure, the pupil time series was averaged across trials, separately for the two conditions (taVNS, sham), resulting in two pupil time series per participant that were used for statistical analyses.

### EEG

#### Data acquisition

The EEG was recorded using a Biosemi Active-Two system (BioSemi) with 64 Ag-AgCl electrodes. To record the vertical and horizontal electrooculogram (EOG), electrodes were placed above and below the left eye and on the outer canthi of both eyes. As described earlier, we also placed electrodes at the back of the earlobe and the auricle to record the stimulation artefact of the taVNS device. EEG, EOG and supporting electrodes were continuously recorded at a sampling rate of 512 Hz and with an electrode impedance < 50 kΩ.

EEG analyses were performed using the EEGLAB toolbox [35], the Fieldtrip toolbox [36], and custom-written routines provided by Sharon and colleagues. EEG data were re-referenced to electrode Cz and high-pass filtered to exclude frequencies below 0.1 Hz. Continuous data were segmented to 15-s epochs (−5 to +10 s) around stimulation onset. Epochs were detrended linearly and notch-filtered at 50 Hz. Then the epoched data were visually inspected to confirm all sham and taVNS trials contained a 25-Hz stimulation artifact. Trials without the artifact (9.4 ± 2.4%) were excluded from all analyses. To focus on the alpha frequency band, we applied a third-order two-pass Butterworth band-pass filter to exclude frequencies below 5 Hz and above 15 Hz. An additional notch filter at stimulation pulse frequency (25 Hz + harmonics up to 100 Hz) was implemented, to remove possible residual artifact activity. Finally, epochs were excluded whenever activity in a channel exceeded ±100 μV in the automatic iterative process specified in the original study. On average, this procedure led to the interpolation of 3.5 ± 0.90 channels and the removal of 0.33 ± 0.13 trials.

After preprocessing was completed, the data were re-referenced to the average reference. The mean number of valid trials was 40.1 ± 1.1 (out of 44) in the taVNS condition, and 37.2 ± 2.0 in the sham condition. Data were downsampled to 64 Hz (Nyquist frequency of 32 Hz > maximum alpha frequency of 15 Hz) and transformed to the time-frequency domain using the Morlet wavelet approach (number of cycles = 7; standard deviations of Gaussian kernel = 3). Applied frequencies increased from 5 to 15 Hz in 31 equally spaced steps. Taken together, this resulted in a frequency resolution of 0.33 Hz and a temporal resolution of 15.6 ms.

To identify participants’ resting-state alpha topography and frequency spectrum, we first extracted data from the breaks between blocks. These were segmented into 5-s epochs, so that there was an overlap of 1 s with both the preceding and subsequent epoch. The obtained epochs underwent the same preprocessing as the stimulation trials and were subsequently reduced to 3-s epochs (discarding the overlap) to avoid filtering artifacts at the edges. This resulted in an average of 34.8 ± 3.6 epochs per participant. Next, we converted the epochs to the time-frequency domain and used them to identify participants’ alpha topography and frequency spectrum using the PARAFAC method [37], as implemented in the N-way toolbox [38]. In line with Sharon et al. [31], we set non-negativity as the constraint for each dimension. The number of components was determined using the core consistency diagnostic (CCD), using a minimal CCD value of 55% as the guiding principle [39]. The CCD amounted to an average of 1.88 ± 0.14 components per participant. Please refer to Sharon et al. [31] for a more detailed depiction of the procedure.

After determination of the optimal number of components, we visually selected the alpha components based on the topographical and frequency distributions, separately for each participant. These were used as weights to be multiplied by the spectrum of all channels. This procedure results in a single channel which represents the weighted activity for both space and frequency. We then applied baseline correction for each trial, by subtracting the mean activity in a time window of -4 to 0 s before stimulation onset.

In line with Sharon et al. [31], we conducted two more analyses on the stimulation data. First, we investigated the time-frequency changes irrespective of the spatial dimension. Therefore, we multiplied the data only with the alpha topographies derived from the PARAFAC decomposition, but ignored the frequency distribution. This allowed us to plot the whole spectrogram at 5-15 Hz as percentage change relative to the same baseline (−4 to 0 s). Second, we investigated the spatial patterns irrespective of frequency distributions. Therefore, we multiplied the data only with the alpha frequency distribution derived from the PARAFAC decomposition, but ignored the topographies. This enabled us to plot the topographical changes relative to the same baseline (−4 to 0 s). Data from both analyses was submitted to cluster-based permutation testing.

### Statistical analysis

Pupil data and data from subjective questionnaires were analysed using Python 3. To estimate the potential effects of taVNS on pupil dilation and subjective feelings of stimulation, we carried out Wilcoxon signed-rank tests using the function *scipy*.*stats*.*wilcoxon*. For analyzing the pupil time series, non-parametric *t*-tests were carried out using *neurotools*.*stats*.*permtest_rel*. Cluster-based permutation analyses for the EEG data were performed using Fieldtrip’s ft_freqstatistics function. We used the Monte-Carlo method and a dependent-samples *t* statistic with 10,000 permutations. Alpha level was set to 0.05 after FDR cluster correction. The time window for the cluster-based permutation analysis was -1 to 6 s. To quantify the effect of taVNS on pupil dilation and alpha power (extracted from the period in which Sharon and colleagues found significant effects of taVNS), we carried out a Bayesian paired samples *t-*tests (using R version 4.0.3, library: *BayesFactor*). We used the default Cauchy prior r = 0.707 in JASP v0.9 [40]. All correlations reported are Spearman’s rank correlation coefficients. Throughout the paper, data are expressed as the mean ± standard error of the mean (SEM).

## Results

Stimulation intensity and participants’ subjective ratings of stimulation are presented in Figure 2. We found no significant differences between the taVNS and sham conditions in levels of subjective averseness (i.e., pain, irritation, mood, alertness; *p*s > .05). Individual stimulation intensities were higher for the sham condition (*M*_*sham*_ = 3.0 ± 1.1 mA) than for the taVNS condition (*M*_*taVNS*_ = 2.3 ± 1.3 mA, *p* = .003), in line with Sharon et al. [31].

**Figure 2.**
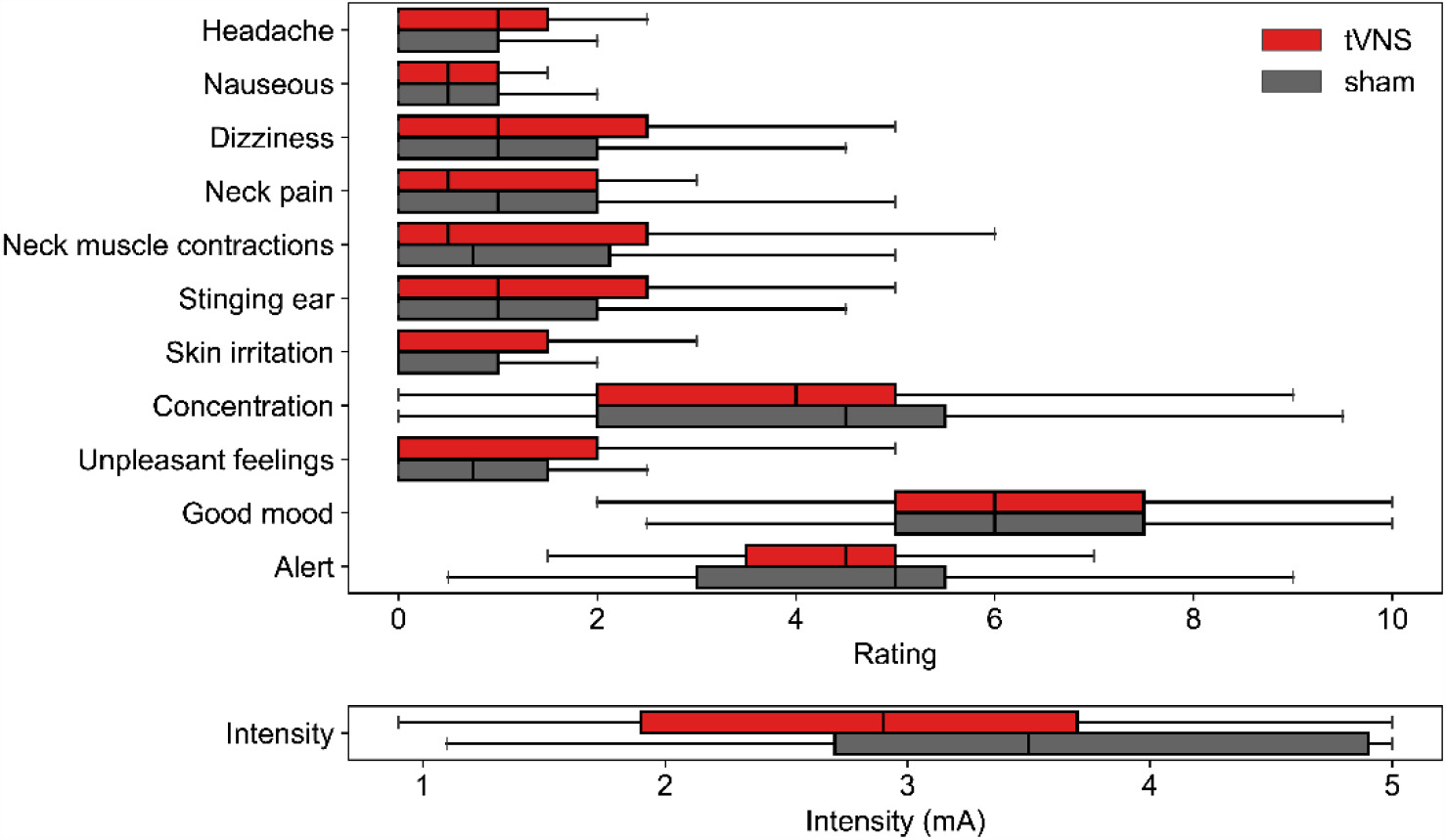
Box plots indicating participants’ subjective ratings of the stimulation, and the objective stimulation intensity levels. Only objective intensity level differed between taVNS and sham.

### Pupillometry

taVNS led to a significant increase in pupil dilation compared to sham stimulation (Figure 3A). Pupil size reached a maximum of 5.6 ± 1.6% above baseline 4.2 s after taVNS onset, in comparison to sham stimulation which peaked at 3.8 ± 1.6% after 4.1 s. Pupil dilation was significantly stronger following taVNS between 2.88 s and 4.90 s (*p*s < .05, two-sided Wilcoxon signed-rank test, FDR-corrected across all timepoints). To determine the replicability of the effect found by Sharon et al. [31], we extracted the average pupil dilation between the two time points in which taVNS led to significantly larger pupil dilation compared to sham in their report (i.e., 2.88 to 5.96 s). Average pupil size was significantly larger for taVNS compared to sham (*p* = .009; Figure 3B). Furthermore, a Bayesian two-sided *t*-test revealed moderate evidence in favor of this effect of taVNS on pupil dilation (BF_10_ = 7.45).

**Figure 3.**
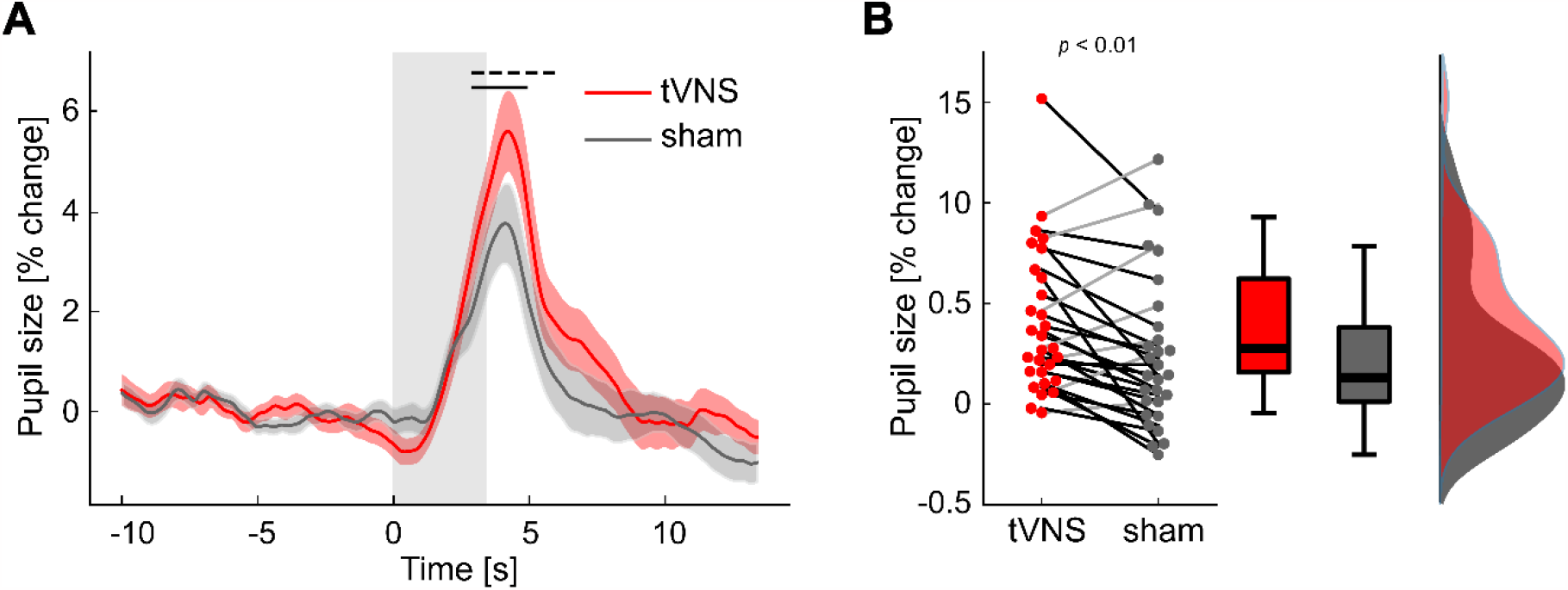
Stronger pupil dilation following taVNS stimulation compared to sham. A) Grand-average pupil dilation waveforms (% change) relative to the 10 s before stimulation onset. Shaded red and grey areas indicate ± SEM. Grey-shaded rectangle indicates the period of stimulation (3.4 s). The horizontal black lines indicate the time period for which pupil dilation differed between taVNS and sham stimulation in the current study (solid line) and for [31] (dashed line). B) Individual data points, box plots and density plots indicating average pupil dilation during the period with a significant taVNS effect in Sharon et al. [31]. Black lines depict participants showing the expected effect (taVNS > sham, *n* = 22), light grey lines depict those who show the opposite effect (sham > taVNS, *n* = 7).

Eye gaze variability and blink rates did not differ between conditions (*p*s > 0.51). Indeed, our key findings remained robust and significant in control analyses in which we either (1) removed pupil data samples where eye gaze in the x or y direction was more than 3 SD away from the mean gaze co-ordinates (i.e., the position of the fixation cross); or (2) did not interpolate over blinks but removed invalid samples from the time series. We also correlated the difference between conditions (taVNS – sham) in objective stimulation intensity and the corresponding difference in evoked pupil response and found no correlation (*R* = 0.18, *p* = .35). However, correlating pupil response and stimulation intensity separately for each condition, we found a positive correlation for taVNS (*R* = 0.39, *p* = .035) but not for sham (*R* = -0.15, *p* = .44). These exploratory, uncorrected, tests reveal tentative evidence that objective stimulation intensity mediates the effect of stimulation on pupil size for taVNS, but not for sham.

### EEG

To identify individual frequency and topography profiles of alpha oscillations for each participant, we employed PARAFAC decomposition to the unbiased break data. In line with Sharon et al. [31], we found individual alpha frequencies between 7 and 13 Hz and an expected occipital topography (Figure 4A). Next, we used these individually derived profiles to obtain weighted average data for the frequency and spatial domain during stimulation trials. Contrary to the original study, we did not find an attenuation of alpha power during taVNS compared to sham in a 4-s time window after stimulation onset (*p* = .89, BF_10_ = 0.23; Figure 4B,C). On a descriptive level, electrical stimulation even *increased* weighted average alpha power, although this effect did not reach significance for taVNS (105 ± 3%, *p* = .12, BF_10_ = 0.42) or sham (104 ± 3%, *p* = .23, BF_10_ = 0.65).

**Figure 4.**
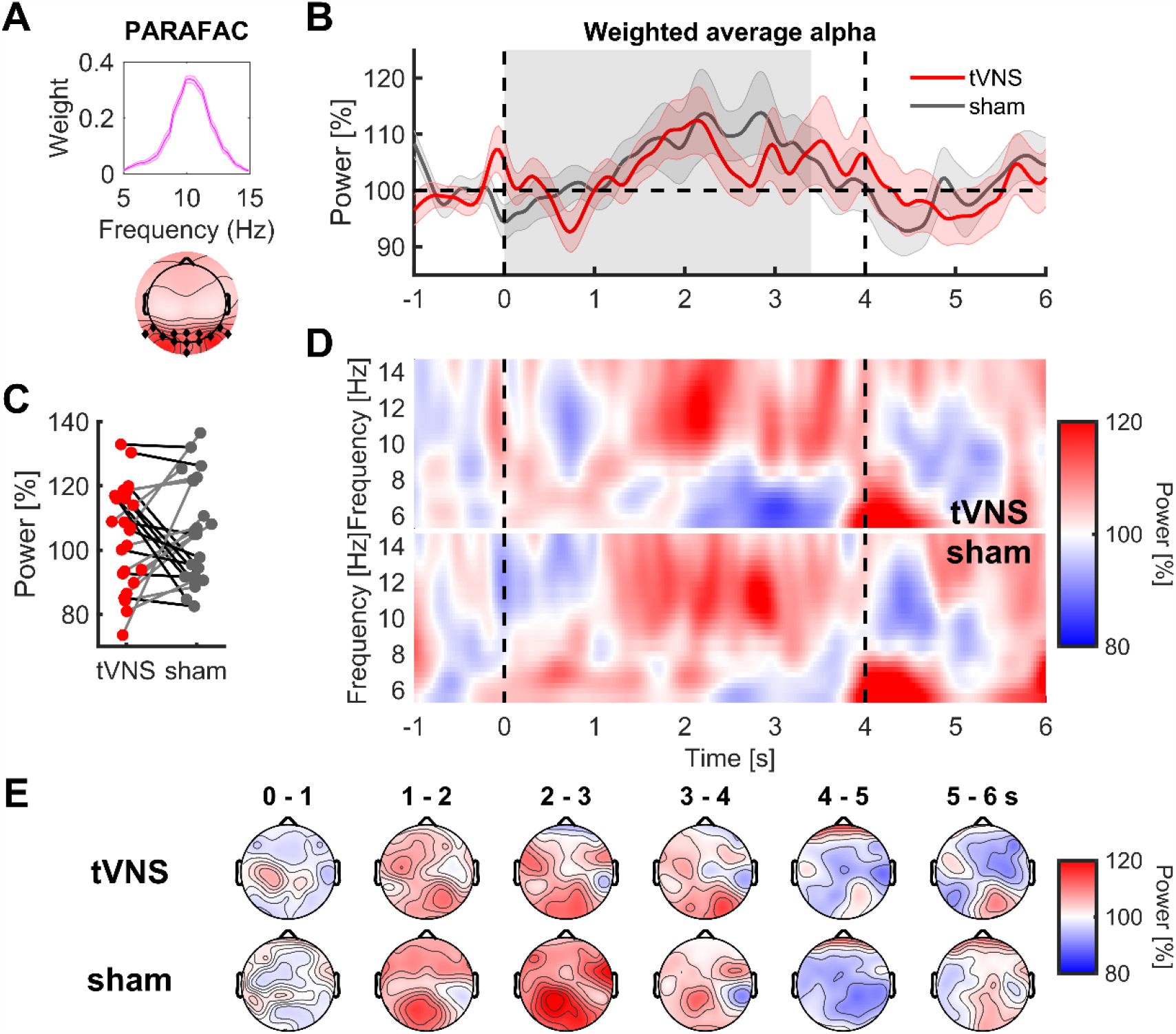
No difference between taVNS and sham in occipital alpha power. A) Median alpha component spectral and spatial profile, as derived from the unbiased break data. B) Grand-average alpha power waveforms, using a weighted average of spectral and spatial profiles in panel A. Dashed vertical lines indicate the time window for the test of significance between taVNS and sham. The grey area denotes the time window of active stimulation (3.4 s). C) Individual data points for the weighted average alpha power in the time window between 0 and 4 s. Grey lines indicate participants with lower alpha for taVNS compared to sham (12 out of 24). Black lines indicate the reversed pattern. D) Mean induced spectrograms, using the weighting of the spatial profile in panel A. E) Mean topographies, using the weighting of the spectral profile in panel A.

Contrary to the original work of Sharon et al. [31], we did not find a significant correlation between the difference in weighted average alpha power (taVNS vs. sham) and the difference in stimulation intensity (*R* = -0.04, *p* = .86). Correlating alpha power and stimulation intensity separately for each condition, we also did not find a link for taVNS (*R* = -0.21, *p* = .33) and sham (*R* = -0.30, *p* = .15).

Following the original study, we also carried out follow-up cluster-based permutation analyses, separately for the time-frequency data (Figure 4D; weighted average over spatial domain) and temporospatial data (Figure 4E; weighted average over frequency domain). In line with our previous results, we did not identify significant clusters of differences between stimulation conditions for the time-frequency data (all *p*s > 0.44) and the temporospatial data (all *p*s > 0.44).

## Discussion

Establishing effective biomarkers of taVNS that are consistent with central noradrenergic activity is relevant for our understanding of how taVNS can be used to study basic cognitive functions and improve clinical applications. We found, in line with Sharon et al. [31], that short taVNS pulses induced a transient pupil response that was larger than that induced by sham stimulation. However, we did not replicate the effects of taVNS on EEG alpha activity. Together, our results contribute to the ongoing research on taVNS by showing that taVNS can alter pupil size, a physiological marker associated with central noradrenergic [41] (and cholinergic [26]) activity. However, it raises questions about the replicability of a taVNS effect on EEG alpha activity.

In accordance with Sharon et al. [31], we found stronger pupil dilation following short bursts of taVNS compared to sham stimulation (BF_10_ = 7.45). This finding is inconsistent with other taVNS studies which used stimulation epochs on the order of minutes [11,19,27–30,42], in some cases with a 30-sec on/30-sec off rhythm. Sharon et al. [31] were the first to assess the effects of short-term taVNS. Since then, others have reported increased pupil dilation following brief (600-ms) taVNS pulse trains, especially when stimulation was applied to the external ear canal [43], although this is not the optimal location for auricular tVNS [44]. The use of short bursts of stimulation could be crucial to trigger phasic, stimulus-evoked noradrenergic responses, as indexed by pupil dilation. For now, there is no evidence that taVNS can modulate *baseline* pupil size.

In addition, Sharon et al. [31] and we used a stimulation intensity level that was individually tailored to be just below painful, while several previous taVNS studies applied substantially lower intensity levels (e.g. 0.5mA) [11,19,28] without catering to individual sensitivity thresholds [30]. Low stimulation intensities may not be capable of activating the vagus nerve afferent fibers near the ear [45]. Indeed, two taVNS studies found parametric effects of stimulation intensity on pupil dilation, which nicely scaled with pulse amplitude [43,46]. These findings, together with evidence from rodents [26], suggest that stronger intensities are more likely to engage the vagus nerve and induce pupil dilation. Future taVNS studies must therefore ensure sufficient intensityꟷwell above the perceptual threshold and below the pain threshold.

We did not replicate the finding of Sharon et al. [31] that short-lasting taVNS transiently attenuated EEG alpha activity; our data were four times more likely under the null hypothesis. This discrepancy is reflective of the state of the literature on VNS and alpha power as measured over the scalp. Little is known about the effect of invasive VNS on alpha power, with the exception of a single-case study that found a significant reduction in alpha power, not during but immediately after 30 seconds of stimulation [47]. Lewine et al. [17] found that 90 seconds of taVNS induced alpha attenuation, but this study used cervical instead of auricular tVNS and involved only eight healthy participants. Ricci et al. [48] found no effect of long-term taVNS on alpha power, also with a limited sample size (n=8). To further complicate matters, Konjusha et al. [49] reported that long-term taVNS in healthy participants modulated alpha power, but in a location-dependent manner: taVNS increased alpha activity over superior parietal regions, but attenuated alpha over middle and superior frontal regions. Finally, Chen et al. [50] found that long-term taVNS attenuated alpha power, but only in the higher alpha-frequency range (∼11 Hz) and in an eyes-closed (but not eyes-open) condition. Furthermore, this effect was largely driven by a difference in alpha activity between the taVNS group and the control group *before* the intervention. Overall, our findings suggest that further research is needed, with larger participant samples and parametric variation of stimulation parameters, to understand the effects of taVNS on alpha activity and to reconcile the mixed findings in the literature.

Finally, it is important to mention a limitation of the current study. Whereas Sharon et al. [31] examined only male participants, we tested mainly female participants. Although this difference could potentially account for the discrepancy between our EEG results, the limited available evidence suggests that female animals [10] and human participants [51] show larger effects of VNS than males, which is inconsistent with our null finding in a group of mainly female participants.

## Data and code availability statement

Preprocessing and analyses code can be found here: https://github.com/bethlloyd/peeg_tvns. Raw pupil time series data and EEG time series data will be made available upon publication.

## Acknowledgement

We thank Omer Sharon for his support and transparency while we carried out this replication study. Omer generously provided scripts to validate our analyses and contributed insightful comments on methodological issues. Moreover, we thank Elio Sjak-Shie for technical support with the robot. We also thank our thesis students and interns who helped with piloting and data collection.

This work was supported by the Netherlands Organization for Scientific Research (grant No. VI.C.181.032).

## Contributions

Conceptualization: S.N., B.L., F.W.; Robot setup and programming: R.d.K, B.L., F.W.; Data collection and curation: B.L., F.W.; Analysis - behavior: B.L.; Analysis - pupil: B.L.; Analysis - EEG: F.W.; Visualization: B.L., F.W.; Writing - original draft: B.L.; Writing - review and editing: S.N., F.W., B.L., R.d.K.; Supervision: S.N.

## Footnotes

1 In case of a discrepancy between the analysis scripts and the information in the article of Sharon et al. [31], we compared both procedures and found essentially the same results. Here we report the results based on the procedures as reported in the scripts.

